# Synthetic mammalian pattern formation driven by differential diffusivity of Nodal and Lefty

**DOI:** 10.1101/372490

**Authors:** Ryoji Sekine, Tatsuo Shibata, Miki Ebisuya

**Affiliations:** RIKEN Center for Biosystems Dynamics Research (RIKEN BDR) 2-2-3 Minatojima-minamimachi, Chuo-ku, Kobe, 650-0047, Japan

## Abstract

Pattern formation is fundamental for embryonic development. Although synthetic biologists have created several patterns, a synthetic mammalian reaction-diffusion pattern has yet to be realized. TGF-β family proteins Nodal and Lefty have been proposed to meet the conditions for reaction-diffusion patterning: Nodal is a short-range activator that enhances the expression of Nodal and Lefty whereas Lefty acts as a long-range inhibitor against Nodal. However, the pattern forming possibility of the Nodal-Lefty signaling has never been directly tested, and the underlying mechanisms of differential diffusivity of Nodal and Lefty remain unclear. Here, through a combination of synthetic biology and theoretical modeling, we show that a reconstituted minimal network of the Nodal-Lefty signaling spontaneously gives rise to a pattern in mammalian cell culture. Surprisingly, extracellular Nodal was confined underneath the cells as small clusters, resulting in a narrow distribution range compared with Lefty. We further found that the finger 1 domain of the Nodal protein is responsible for its short-range distribution. By transplanting the finger 1 domain of Nodal into Lefty, we converted the originally long-range distribution of Lefty to a short-range one, successfully preventing the pattern formation. These results indicate that the differences in the localization and domain structures between Nodal and Lefty, combined with the activator-inhibitor topology, are sufficient for reaction-diffusion pattern formation in mammalian cells.

## Main

One of the goals of synthetic biology is creating a synthetic tissue to understand natural developmental mechanisms^1-3^, to explore the origin of multicellularity^4^ and to engineer a programmable tissue for therapeutic purposes^5,6^. The first step towards a synthetic tissue is controlling pattern formation, which enables to place different types of cells properly in a tissue. Several synthetic cellular patterns have been reported previously: Ring patterns were created in genetically engineered bacteria that can sense the concentrations of small molecules^7,8^. In mammalian cells, 2D and 3D patterns were created based on engineered cell sorting mechanisms^9,10^. However, there is another pattern formation mechanism that has not been artificially created in mammalian cells despite its importance: the reaction-diffusion (RD) patterning system.

The concept of a self-organizing RD system was first proposed by Alan Turing as a chemical system of interacting and diffusible molecules giving rise to various stable patterns, such as spots and stripes^11-14^. Recent studies have suggested that RD system underlies a number of developmental patterning phenomena, including digit formation in the limb^15,16^, pigmentation on the skin^17^, the formation of hair follicles and feather buds on the skin^18,19^, branching morphogenesis in the lung^20^ and rugae formation in the palate^21^. In the field of synthetic biology, a regular stripe pattern has been created in colonies of engineered bacteria, in which diffusion of the small molecule AHL regulates the motility of the actively-swimming bacteria^22^, and this patterning mechanism can be considered as a non-classic RD system. Very recently, a stochastic Turing pattern has been created in engineered bacteria that have a synthetic network of two diffusible small molecules^23^. However, an RD pattern has not so far been created in eukaryotic cells. Furthermore, although RD patterning in embryonic development is mediated mostly by the interaction of diffusible protein ligands called morphogens rather than by small molecules or cell movement^15,16,18-21^, a protein-based RD patterning system has not so far been created either.

Our goal here thus was to engineer a minimal RD patterning network of protein ligands in mammalian cells. An RD pattern requires a minimum of two diffusible molecules, or signaling pathways, and we chose to employ the well-studied Nodal-Lefty signaling pathway, which regulates mesodermal induction, axis formation and left-right patterning^24-26^. The Nodal-Lefty pathway has been proposed to meet two conditions for a stable RD pattern^14,27^: Firstly, binding of Nodal to its receptor activates the production of both Nodal and Lefty whereas Lefty inhibits the activity of Nodal^24,28,29^. Secondly, the diffusion of Nodal is reported to be slower than Lefty in zebrafish, chick and mouse embryos^27-31^. In other words, Nodal and Lefty may act as a short-range activator and a long-range inhibitor, respectively, and thus satisfy the requirement for a classic Turing pattern proposed by Gierer and Meinherdt^12,13^. It remains undemonstrated, however, whether the Nodal-Lefty signaling can actually produce an RD pattern, as well as how Nodal and Lefty show different diffusivity.

In this study, we have reconstituted a minimal activator-inhibitor circuit of Nodal and Lefty to test if it leads to any pattern formation in mammalian cell culture. We also took advantage of the simple *in vitro* system and investigated the differences in the diffusion mechanisms of Nodal and Lefty.

## Results

### Synthetic pattern formation with an activator-inhibitor circuit

We first created an activator circuit in HEK293 cells, where the activator Nodal induces the expression of Nodal itself (Fig. 1a). Extracellular Nodal is known to bind to the co-receptor, Cryptic or Cripto, as well as to the heterodimeric receptors, Activin receptor type I and II. The activated receptors then activate Smads that form a complex with the transcription factor FoxH1, leading to the transcription of downstream targets^24^. Since HEK293 cells lack some of these essential components to transduce the Nodal signaling^32^, we introduced exogenous Cryptic and FoxH1 to the cells (Supplementary fig. 1). The induction of gene expression in response to Nodal signaling was monitored with an (f2)_7_-luc reporter, a seven-times repeat of a FoxH1-responsive element that regulates the luciferase expression^33^. The exogenous expression of a type II receptor, Acvr2b, further improved the induction rate of the (f2)_7_-luc reporter signal (Supplementary fig. 1). Co-culturing with Nodal-producing cells, instead of the recombinant Nodal proteins, also activated the (f2)_7_-luc reporter cells (Supplementary fig. 2a), showing that secreted Nodal propagates to the neighboring cells. Finally, we added the (f2)_7_-Nodal construct to let the cells both produce and respond to Nodal, completing the positive feedback of the activator circuit (Fig. 1a). When the HEK293 cells engineered with the activator circuit were seeded at a high density (near confluent), the (f2)_7_-luc reporter signal was initially detected only in ~10% of the cells, and those reporter-positive cells were randomly distributed (Fig. 1b, 0 h). Then the reporter-positive cells activated the immediate neighboring cells, and the small domains of the positive cells appeared at around 18 hours. By 42 hours, all the cells became reporter-positive (Fig. 1b,c; Supplementary movie 1). Note that the cells are nearly confluent from the time zero and that the cell proliferation rate is low (the cells divided approximately twice in 70 hours), meaning that the observed signal propagation is not because of the cell proliferation but because of the mutual activation of Nodal signaling among the neighboring cells. The propagated reporter signal lasted until 50-60 hours and started to gradually decline possibly due to the lack of fresh medium.

**Figure 1.**
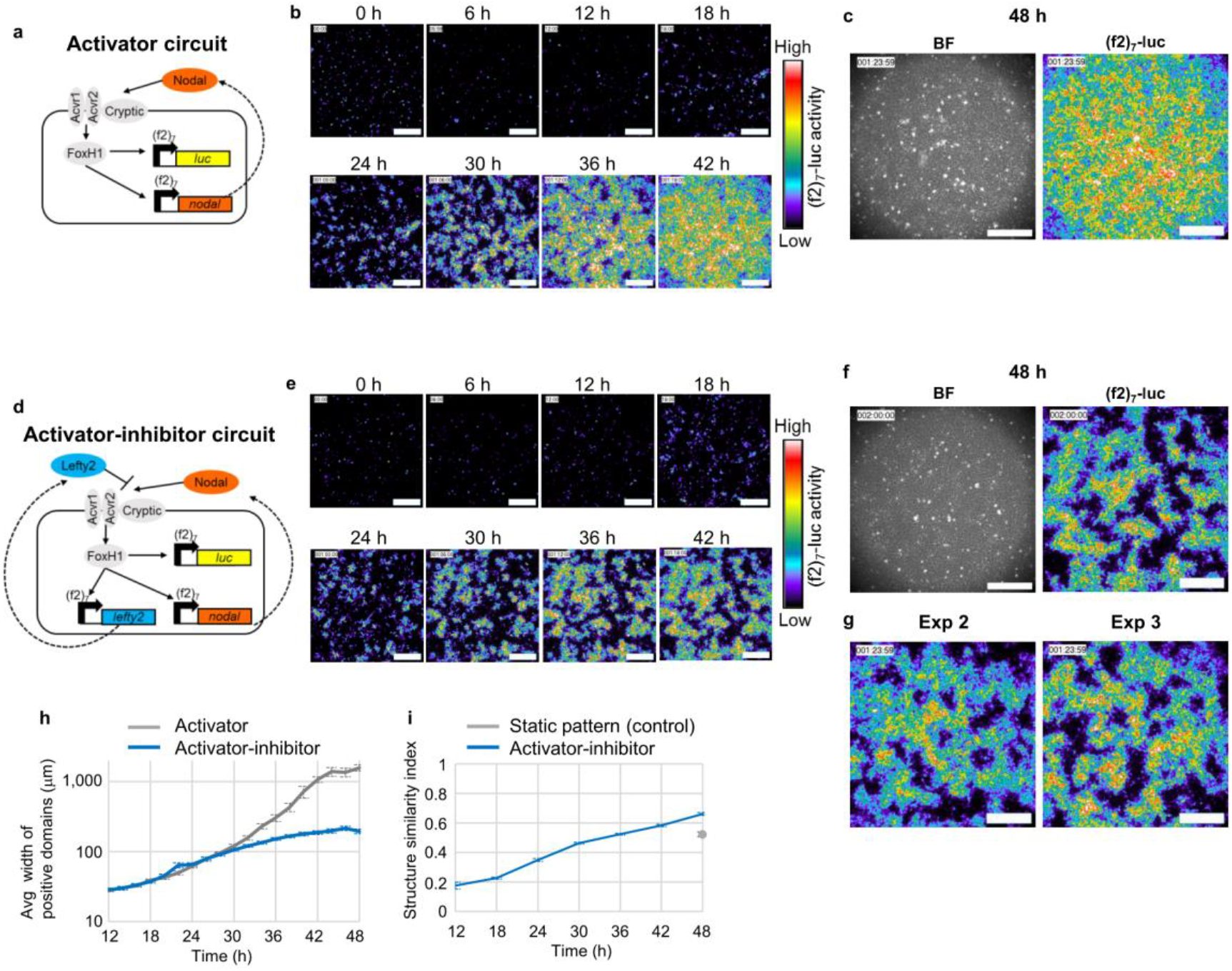
Cells with an activator-inhibitor circuit spontaneously give rise to a pattern. (**a**) The activator circuit. (**b**) Time-lapse imaging of the HEK293 cells engineered with the activator circuit. See also Supplementary movie 1. (**c**) The bright filed and luciferase images of the cells with the activator circuit at 48 hours. (**d**) The activator-inhibitor circuit. (**e**) Time-lapse imaging of the HEK293 cells engineered with the activator-inhibitor circuit. See also Supplementary movie 1. (**f**) The bright filed and luciferase images of the cells with the activator-inhibitor circuit at 48 hours. (**g**) Repeated experiments of (f). (**h**) The width of positive domains was measured at each time point as described in Supplementary fig. 3. (**i**) The structural similarity (SSIM) index between two images at time t and t + 6 hours was calculated as described in Supplementary fig. 3 A. higher index means a more stable pattern. The gray dot at 48 hours indicates the SSIM index of a control static sample, where the cells that constitutively express luciferase were mixed with wild-type cells.

This activator circuit serves as a base for an activator-inhibitor circuit, where Nodal induces the expression of Nodal as well as that of Lefty that inhibits the Nodal signaling (Fig. 1d). We first confirmed that co-culturing with Lefty2-producing cells inhibits the activation of the (f2)_7_-luc reporter cells (Supplementary fig. 2b). Then we introduced the (f2)_7_-Lefty2 construct into the HEK293 cells already engineered with the activator circuit, adding the negative feedback by Lefty (Fig. 1d). The activator-inhibitor circuit initially behaved similarly to the activator circuit: the small domains of reporter-positive cells appeared at around 18 hours and became bigger (Fig. 1e). Then the domain growth slowed down at around 30 hours, and a pattern of clear positive domains and negative domains was formed by 36 hours (Fig. 1e,f; Supplementary movie 1). Note that the reporter-positive cells and negative cells are genetically identical since the cell line was cloned. The pattern was reproducible (Fig. 1g), and the average size of positive domains was 196 ± 15 μm (Fig. 1h; Supplementary fig. 3). The pattern did not change much after 36 hours and kept essentially constant until 60 hours (Fig. 1h,i), even though the entire signal started to decline at 50-60 hours as observed with the activator circuit.

To assess the periodicity of our synthetic pattern, we calculated the spatial correlation of the image (Supplementary fig. 4a-d). The second peak of the radially averaged correlation function indicates the distance over which the pattern repeats itself (i.e., the distance from peak-to-peak of a pattern). While the correlation of the image of the cells with the activator circuit rapidly dropped (Supplementary fig. 4b), that with the activator-inhibitor circuit showed a small second peak at around 400 μm (Supplementary fig. 4d), suggesting a weak periodicity of the pattern. This period was consistent with the sum of the average positive domain width and negative domain width (376 ± 99 μm) (Supplementary fig. 4e). These results indicate that our activator-inhibitor circuit, a minimal network of the Nodal-Lefty signaling, has an ability to make cells form a stable RD pattern with a periodicity of ~400 μm.

### Different distribution ranges of Nodal and Lefty

Why can the synthetic Nodal-Lefty circuit give rise to a pattern? Since a stable RD pattern typically requires a short-range activator and a long-range inhibitor^11-14^, the difference in the diffusion ranges of two diffusible molecules is critical. Although the diffusion of Nodal has been reported to be slower than that of Lefty in zebrafish, chick and mouse embryos^27-31^, our experimental conditions are different from those of the previous studies especially because we culture the cells on a dish as a monolayer with plenty of culture medium. We thus tested whether Nodal and Lefty show different diffusion ranges in our system. To visualize the distribution of Nodal and Lefty, we placed the ligand-producing cells and the receptor cells (i.e., wild-type cells) separately in adjacent areas by using a mold called a culture insert (Fig. 2a). Then the extracellular Nodal and Lefty were exclusively visualized with a split luciferase system called HiBiT (Fig. 2b): The N-terminal bigger half of NanoLuc, Large BiT (LgBiT), is added to the medium but does not enter a cell. Thus, the C-terminal smaller half of NanoLuc, the HiBiT tag, binds to LgBiT to reconstitute a functional luciferase only outside the cell (Fig. 2b). Avoiding intracellular signal this way is crucial since the concentrations of Nodal and Lefty are much higher inside the cell, which easily masks their extracellular distributions. Because Nodal and Lefty are cleaved by proteases to become their mature forms^28,34^, we inserted the HiBiT tag into the middle part of the proteins, at the N-terminus of the mature domains (Fig. 2c,d).

**Figure 2.**
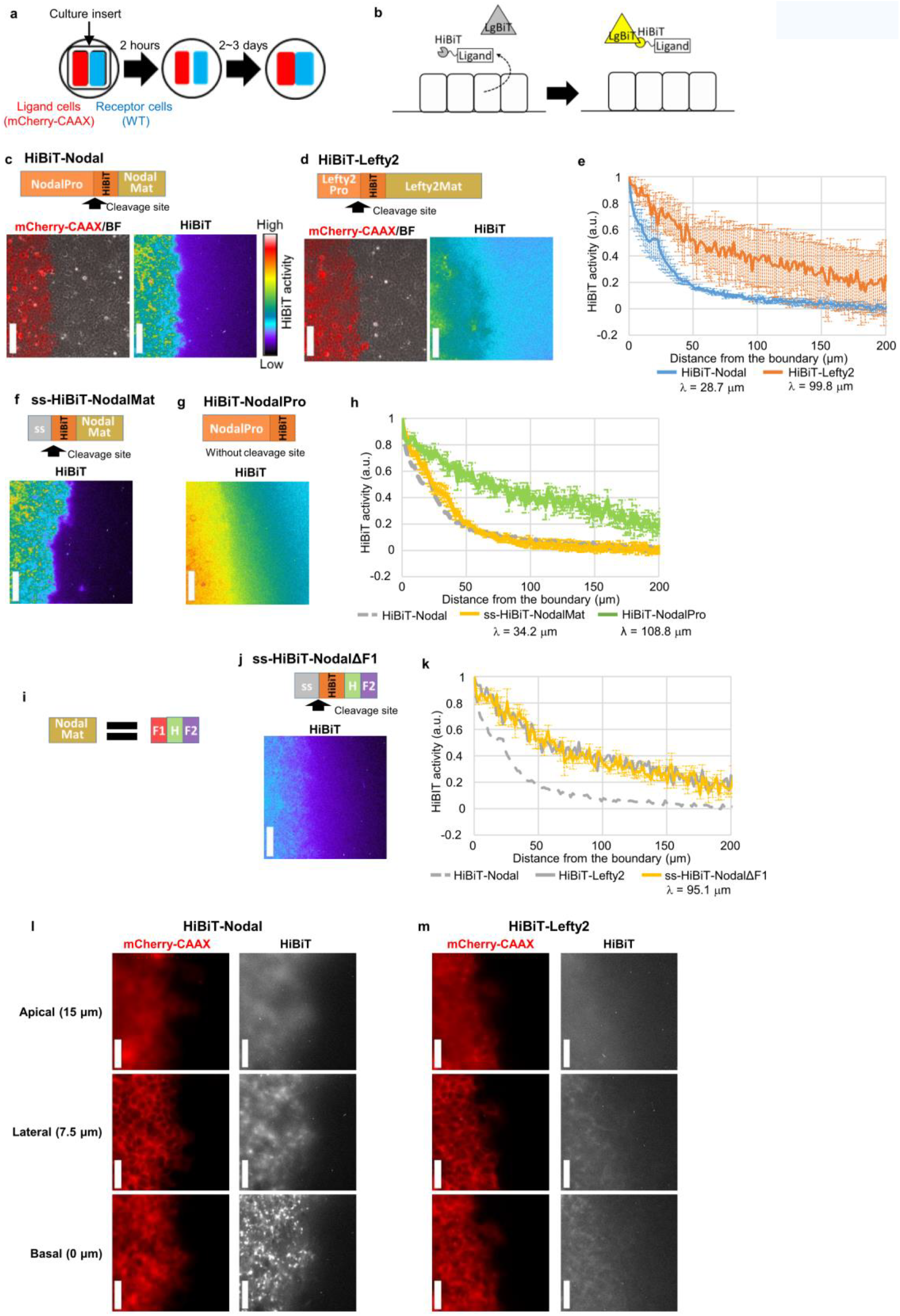
The distribution range of Nodal is shorter than that of Lefty. (**a**) A scheme of the culture insert assay. The ligand-producing cells are labeled with mCherry-CAAX. The ligand cells and the receptor cells (wild-type cells) are cultured separately in the two wells of a culture insert for 2 hours. After the culture insert is removed, the cells fill the cell-free gap, establishing a straight boundary between the two cell types. (**b**) A scheme of the HiBiT system to visualize the extracellular distribution of ligands. The small tag, HiBiT, is fused to the ligand protein, whereas the LgBiT and substrate are added to the medium. The HiBiT and LgBiT reconstitute NanoLuc only outside the cells. (**c**) Top: The HiBiT tag was inserted into the N-terminus of the Nodal mature domain. Bottom left: The boundary of the HiBiT-Nodal-producing cells (ligand cells, labeled with mCherry-CAAX) and receptor cells. The mCherry-CAAX image was merged with the bright field image. Bottom right: The HiBiT-mediated luminescence image showing the extracellular distribution of HiBiT-Nodal. (**d**) Top: The HiBiT tag was inserted into the N-terminus of the Lefty2 mature domain. Bottom left: The boundary of the HiBiT-Lefty2-producing cells and receptor cells. Bottom right: The extracellular distribution of HiBiT-Lefty2. (**e**) Quantified distribution profiles of HiBiT-Nodal and HiBiT-Lefty2. Each distribution was fitted to exp(-x/λ) to estimate the characteristic distance λ. (**f**) Top: A signal sequence (ss) and the HiBiT tag were fused to the N-terminus of the Nodal mature domain. Bottom: The extracellular distribution of ss-HiBiT-NodalMat. (**g**) Top: The HiBiT tag was fused to the C-terminus of the Nodal prodomain. Bottom: The extracellular distribution of HiBiT-NodalPro. (**h**) Quantified distribution profiles of ss-HiBiT-NodalMat and HiBiT-NodalPro. The HiBiT-Nodal distribution shown in (e) is displayed as a control. (**i**) The Nodal mature domain consists of three subdomains: the finger 1 (F1), heel (H) and finger 2 (F2). (**j**) Top: The finger 1 domain was deleted from ss-HiBiT-NodalMat. Bottom: The extracellular distribution of ss-HiBiT-NodalΔF1. (**k**) Quantified distribution profiles of ss-HiBiT-NodalΔF1. The distributions of HiBiT-Nodal and HiBiT-Lefty2 shown in (e) are displayed as a control. (**l, m**) Higher magnification view of HiBiT-Nodal (**l**) and HiBiT-Lefty2 (**m**). The apical, lateral, basal sides were defined as described in Supplementary fig. 6. The mCherry-CAAX images were normalized differently between (l) and (m). Data are means and s.e.m. (n=3). Scale bars: 200 μm (c,d,f,g,j); 50 μm (l,m).

The luminescence signal of HiBiT-Nodal displayed an extremely narrow distribution (Fig. 2c). The Nodal distribution reached at its equilibrium by four hours after the addition of LgBiT and substrates (Supplementary fig. 5a). To compare the distribution ranges of different proteins, we fitted the normalized distribution to a simple exponential function, exp(-x/λ). The characteristic distance λ is the point where the signal drops to 1/e, and λ also represents 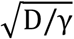, where D is the effective diffusion coefficient and γ is the degradation rate^35,36^. In the case of HiBiT-Nodal, λ = 28.7 μm (Fig. 2e), suggesting that the effective range of Nodal is only one or two cells since the cell size is 10-20 μm. By contrast, HiBiT-Lefty2 displayed much wider distribution where the signal gradually decreased (Fig. 2d), and λ = 99.8 μm (Fig. 2e), indicating that the effective range of Nodal is 3.5-times narrower than that of Lefty2.

The molecular sizes of Nodal and Lefty2 are similar (full-length Nodal: 354 aa; full-length Lefty2: 368 aa; mature Nodal: 110 aa; mature Lefty2: 291 aa) and thus unlikely to be the cause of their different diffusion ranges. We hypothesized the existence of a trap mechanism to confine extracellular Nodal in immediate neighboring cells, and focused on Nodal rather than Lefty. Since the full-length Nodal protein comprises the mature domain and prodomain (Fig. 2c), we examined which domain is responsible for the narrow distribution. Whereas the mature domain of Nodal displayed a narrow distribution just like the full-length Nodal (Fig. 2f), the prodomain displayed a wider distribution just like Lefty2 (Fig. 2g,h). The Nodal mature domain further comprises three subdomains^37^: the finger 1 domain, heel domain and finger 2 domain (Fig. 2i). Deleting the finger 1 domain from the Nodal mature protein made the distribution wider (Fig. 2j,k), indicating that the finger 1 subdomain of Nodal is responsible for its narrow distribution.

### Extracellular Nodal localizes underneath the cells

We further investigated how the finger 1 domain limits the distribution range of Nodal. While a previous study has suggested that binding of Nodal to the Acvr2b receptor slows down the Nodal diffusion in zebrafish^38^, the overexpression or deletion of Acvr2b in our system did not change the Nodal distribution range (Supplementary fig. 5b). Then we checked the localization of extracellular Nodal, noticing that the HiBiT-Nodal signal was in focus at the basal side of cells but out of focus at the lateral or apical side (Fig. 2l). The basal side was judged with the dense structure of cell membrane, and the lateral and apical sides were defined as the points 7.5 μm and 15 μm above the basal side, respectively (Supplementary fig. 6). Consistent with this observation, extracellular Nodal is suggested to localize underneath the cells even in mouse embryos^39^. We also noticed that the HiBiT-Nodal near the basal side formed small clusters (Fig. 2l). By contrast, the HiBiT-Lefty2 signal was blurry both at the basal and apical sides (Fig. 2m). These results suggest that extracellular Nodal is confined in the space between the cells and the culture dish as clusters, which may be the cause of the narrow distribution of Nodal. To test the involvement of cell-dish attachment, we treated the cells with an Actin inhibitor Cytochalasin B that disrupts the focal adhesions, finding that the strong signal of HiBiT-Nodal clusters near the basal side of cells disappeared quickly (Supplementary fig. 7, 3h).

### Mathematical models of the pattern forming circuit

To understand the patterning mechanism of our activator-inhibitor circuit in more detail, we constructed simple mathematical RD models (Fig. 3a-j; Methods). Two mechanisms have been reported regarding how Lefty inhibits the Nodal signaling: Lefty competes with Nodal for the co-receptor and receptors^28,40^ (Fig. 3a), or Lefty directly binds to and then inhibits Nodal^40^ (Fig. 3e). We thus constructed two types of model by using parameters we measured or estimated (Supplementary fig. 8): the “competitive inhibition model” and the “competitive inhibition + direct inhibition model”. Both models gave rise to patterns comprising positive domains and negative domains when the parameters were in the right ranges (Fig 3b,f). The patterns resulting from the competitive inhibition model were highly periodic (Fig. 3d), and the patterning parameter range was almost identical with the parameter range that satisfied Turing instability^13^ (compare Fig. 3c with 3b), the condition for Turing pattern formation, meaning that these patterns are classic Turing patterns. However, Turing patterns are not the only type of RD system that can perform spatial patterning. The patterns resulting from the competitive inhibition + direct inhibition model were less periodic (Fig. 3h,i), and the Turing instability condition was not satisfied in all the parameter regions tested with this model (Fig. 3g). This non-Turing patterning mechanism is essentially the same as the formation of “solitary localized structures”^41,42^ caused by an excitable or bistable system combined with a rapidly diffusing inhibitor: the positive domains are formed by short-range self-activation initially, and the propagation of domains are stopped by long-range inhibition in the later stage. Thus, we named the less-periodic patterns as “solitary patterns”. Since the periodicity of the actual cell pattern resulting from our activator-inhibitor circuit was not very clear (Supplementary fig. 4d), our synthetic RD pattern may be a solitary pattern rather than a Turing pattern.

**Figure 3.**
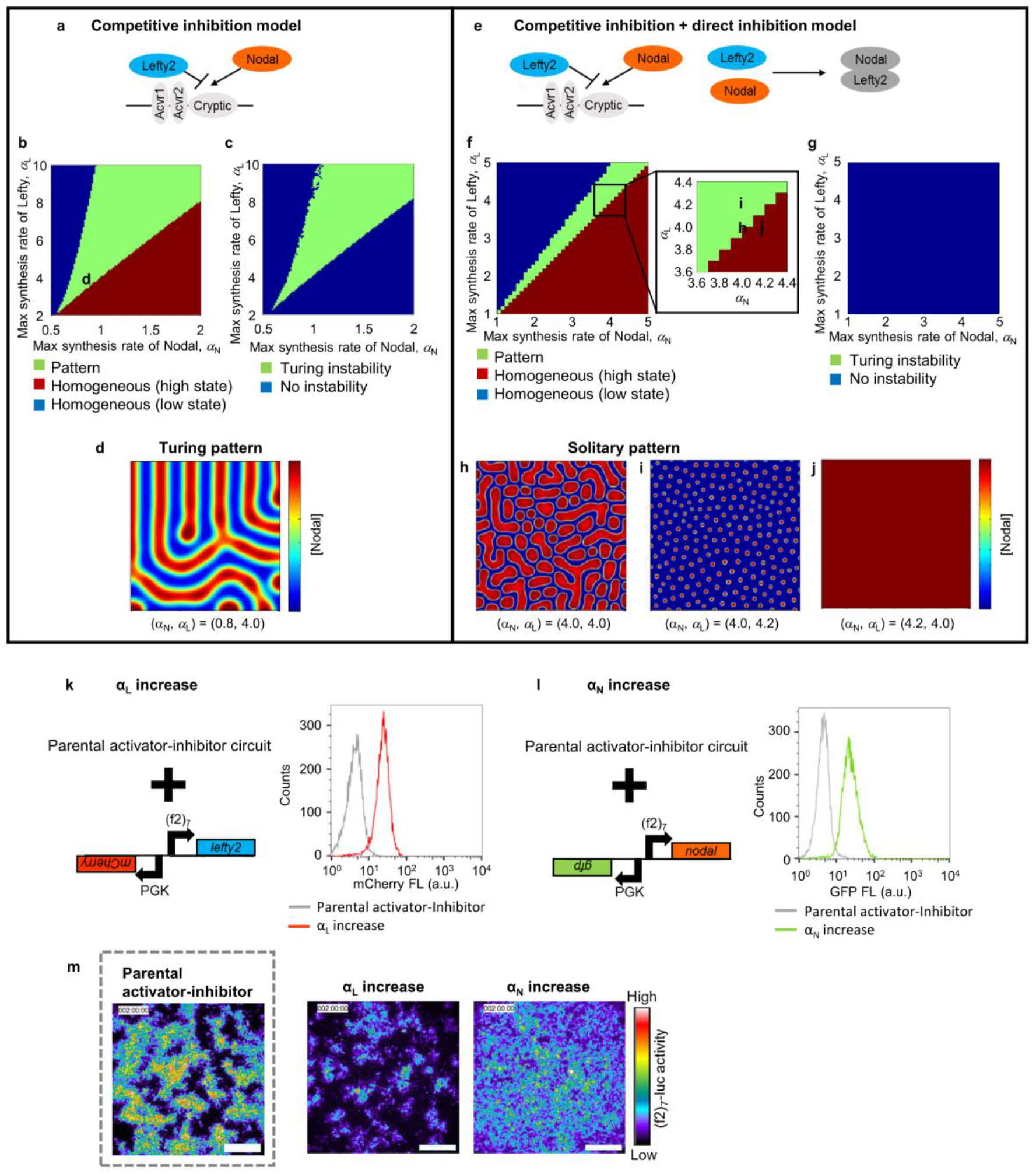
Mathematical models of the activator-inhibitor circuit. (**a-d**) The competitive inhibition model. (**a**) A scheme of competitive inhibition. (**b**) The parameter region for pattern formation (green). The competitive inhibition model was simulated in one dimension. See also Methods. The parameter combination used in (d) is indicated by the character d. (**c**) The parameter region that meets the Turing instability condition (green). (**d**) Two-dimensional simulation with (α_N_, α_L_) = (0.8, 4.0). A typical Turing pattern was formed with this model and parameter set. The parameter combination used in (d) are indicated in (b). (**e-j**) The competitive inhibition + direct inhibition model. (**e**) A scheme of competitive inhibition and direction inhibition. (**f**) The parameter region for pattern formation (green). The competitive inhibition + direct inhibition model was simulated in one dimension. The parameter combinations used in (h-j) are indicated in the inset. (**g**) The Turing instability condition was not satisfied in the entire parameter regions tested. (**h-j**) Two-dimensional simulation with different combinations of the parameters αN and αL. Solitary patterns, not Turing patterns, were formed with this model and parameter range. (**k**) Left: To increase αL, a construct containing (f2)_7_-Lefty2 and PGK-mCherry was added to the parental activator-inhibitor circuit shown in Fig. 1d. Right: FACS plot for the PGK-mCherry intensity, confirming the introduction of the construct. (**l**) Left: To increase αN, a construct containing (f2)_7_-Nodal and PGK-GFP was added to the parental activator-inhibitor circuit. Right: FACS plot for the PGK-GFP intensity. (**m**) Patterns resulting from the activator-inhibitor circuit with the increased α_N_ or α_L_. The pattern of the parental activator-inhibitor cells shown in Fig. 1f is displayed as a control. Scale bars, 400 μm.

An important step in theoretical modeling is to alter parameters in the model, and then test the observed predictions experimentally. We thus attempted to alter the maximum synthesis rate of the activator-inhibitor circuit both in simulation and living cells. Our model predicted that increasing the maximum synthesis rate of Nodal or Lefty should change the ratio of the positive domains to negative domains in the resulting patterns (Fig. 3h-j). To experimentally increase the maximum synthesis rates, we introduced the extra copies of (f2)_7_-Lefty2 or (f2)_7_-Nodal to the cells already engineered with the activator-inhibitor circuit (Fig. 3k,l). The constitutive expression of PGK-mCherry or PGK-GFP from the same construct as (f2)_7_-Lefty2 or (f2)_7_-Nodal was used as a marker of the increased copy numbers (Fig. 3k,l; FACS plots). As predicted, increasing the maximum synthesis rate (i.e., the copy number) of Nodal made the cells homogeneously positive, whereas that of Lefty expanded the area of negative domains (compare Fig. 3m with Fig. 3h-j).

### Manipulating the diffusion coefficient of Lefty

We further altered another important parameter, the diffusion coefficient of the activator-inhibitor circuit. Our model predicted that no pattern should be formed, irrespective of a Turing pattern or a solitary pattern, if the diffusion coefficients of Nodal and Lefty are the same (Compare Fig. 4a with Fig. 3f). To experimentally test this prediction, we fused the finger 1 domain of Nodal to Lefty2 (Fig. 4b). As expected, the HiBiT-F1-Lefty2 displayed a narrow distribution just like Nodal (Fig. 4b,c). The HiBiT-F1-Lefty2 also localized near the basal side of cells and formed small clusters, just like Nodal (compare Fig. 4d with Fig. 2l,m). These results show that the Nodal finger 1 domain is indeed important for the narrow distribution and able to make the distribution of Lefty2 narrow upon transplantation.

**Figure 4.**
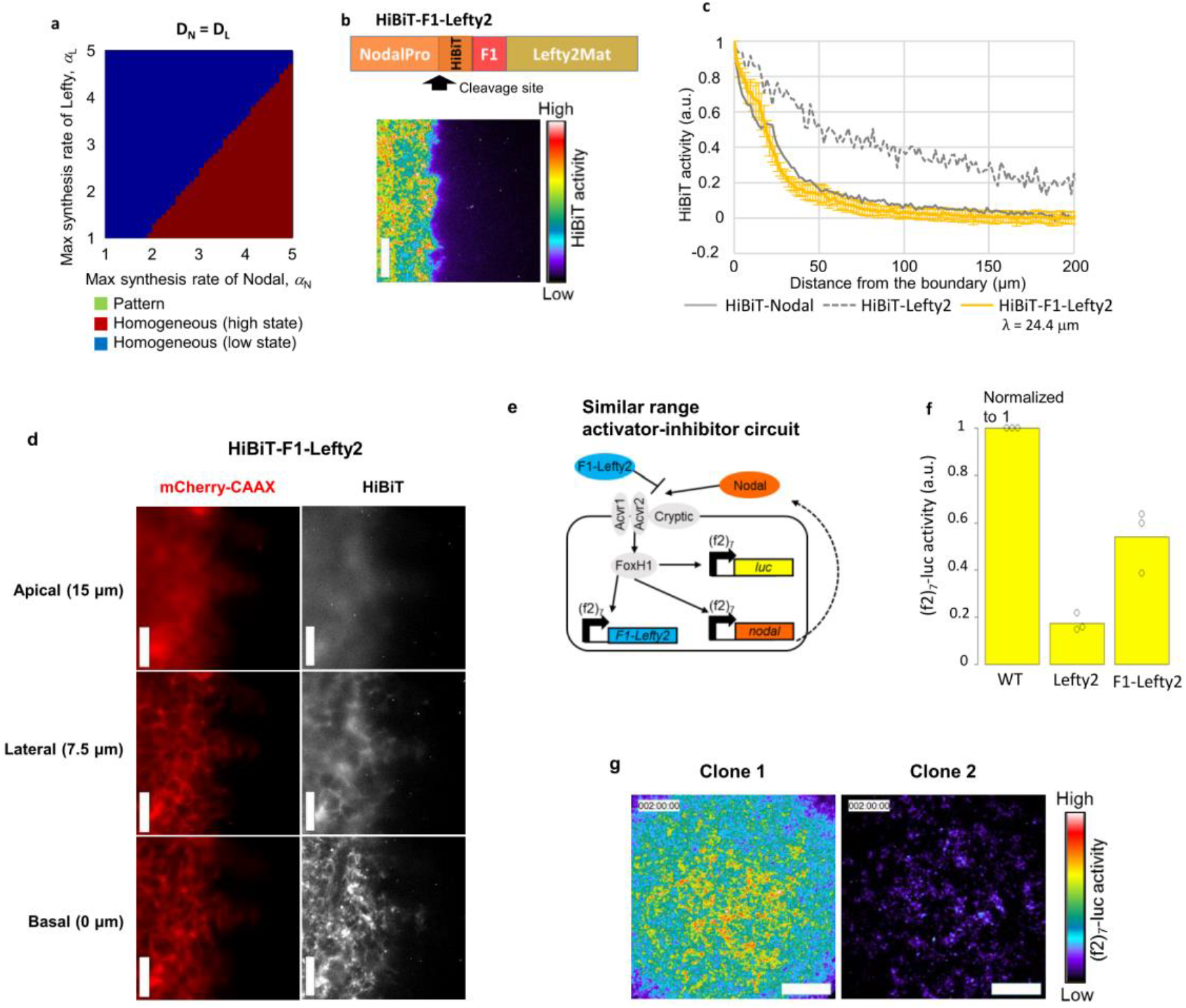
Different diffusivity of Nodal and Lefty is crucial for the pattern formation. (**a**) The parameter region for pattern formation does not exist with the same diffusion coefficients for Nodal and Lefty (D_N_ = DL = 1.96 μm^2^/min). The competitive inhibition + direct inhibition model was simulated in one dimension. (**b**) Top: The finger 1 domain of Nodal was fused to Lefty2. Bottom: The culture insert assay showing the extracellular distribution of HiBiT-F1-Lefty2. (**c**) Quantified distribution profile of HiBiT-F1-Lefty2. The distributions of HiBiT-Nodal and HiBiT-Lefty2 shown in Fig. 2e are displayed as a control. Data are means and s.e.m. (n=3). (**d**) Higher magnification view of HiBiT-F1-Lefty2. (**e**) The similar range activator-inhibitor circuit. F1-Lefty2 was used instead of Lefty2 in the activator-inhibitor circuit to make the diffusion coefficients of Nodal and Lefty similar. (**f**) Inhibition activity of F1-Lefty2. Wild-type cells, Lefty2-producing cells or F1-Lefty2-producing cells were co-cultured with the cells engineered with the activator circuit, and the (f2)7-luc activities were measured. Data are means and individual points (n=3). (**g**) The image of two representative clones of the cells engineered with the similar range activator-inhibitor circuit at 48 hours. Scale bars, 200 μm (b); 50 μm (d); 400 μm (g).

Then we created an activator-inhibitor circuit with F1-Lefty2, instead of wild-type Lefty2, and named the new circuit as a “similar range activator-inhibitor circuit” (Fig. 4e). F1-Lefty2 inhibited the Nodal signaling although its inhibitor activity was a little weaker than that of wild-type Lefty2 (Fig. 4f). When the (f2)_7_-F1-Lefty2 construct was added to the activator circuit, the engineered cells did not show a pattern but displayed an almost homogeneous image (Fig. 4g), and the spatial correlation dropped rapidly without showing a second peak (Supplementary fig. 9). These results confirmed our prediction that different diffusion ranges of Nodal and Lefty are crucial for the pattern formation through our activator-inhibitor circuit.

## Discussion

We have created here, to our knowledge, the first synthetic RD pattern in mammalian cells, that is verified by comparison of various experimental perturbations to a theoretical model (i.e., changing the diffusion constants and maximum expression levels). In doing so, we show that the pattern formation through our synthetic circuit is driven by the different diffusion ranges of Nodal and Lefty, which are influenced by the Nodal finger 1 domain and the confinement of Nodal underneath the cells.

Extracellular Nodal localized underneath the basal side of cells. Although the mechanism for this localization remains unclear, one possibility is that Nodal is trapped by protein complexes that exist between the cells and a dish, such as the extracellular matrix (ECM) and adhesion complex. In fact, Nodal is reported to interact with sulfated glycosaminoglycans that localize to the basement-membrane like structure underneath the cells in mouse embryos^39^. Since Nodal and Lefty are known to display different diffusivity in developing mouse, chick and zebrafish embryos, it will be interesting to examine whether the Nodal localization as well as the finger 1 domain affect the distribution range of Nodal in those embryos. The localization underneath the cells enabled Nodal to form a steep gradient in our cell culture even with plenty of culture medium. Without a proper trapping mechanism, the free diffusion in the medium should be too fast for Nodal to form any distribution. Recent studies show the gradient distribution formation of other morphogens in cell culture^43^, suggesting that localization underneath the cells may be a common trapping mechanism.

Nodal displayed a 3.5-times shorter distribution range than Lefty2 in our measurements. If the degradation rates are similar between Nodal and Lefty2, the effective diffusion coefficient of Nodal should be 12-times smaller than that of Lefty2. According to our measurements, the degradation of Lefty2 is actually 2.4-times faster than that of Nodal (Supplementary fig. 8b), suggesting that the effective diffusion coefficient of Nodal is 29-times smaller than that of Lefty2, which is comparable to the value reported in zebrafish^27^. Direct measurements of the diffusion coefficients in our system will be necessary to verify these numbers even though our attempts for FRAP analysis were not successful due to too weak signal of fluorescent fusion Nodal and Lefty. In any case, sufficiently large difference in the diffusivity of Nodal and Lefty was proven critical for our pattern formation.

The cells engineered with our activator-inhibitor circuit gave rise to a pattern, which we believe is the first mammalian example of a synthetic RD pattern. Our mathematical models suggested two possible RD systems: a Turing pattern and a solitary pattern. Both patterns require the positive feedback of a short-range activator and the negative feedback of a long-range inhibitor. The uniform stationary state is absolutely unstable in Turing periodic patterns, whereas it is stable in solitary spot patterns. Therefore, a solitary pattern can be initiated at the position where sufficiently strong local perturbation is applied. By contrast, a Turing pattern might appear without any significant perturbation. Although our synthetic cell pattern looks more similar to a solitary pattern compared with a Turing pattern, further experiments are needed to distinguish the two possibilities.

While pattern formation is fundamental for embryonic development, investigating the underlying molecular mechanisms in complex living tissues is often difficult. A simple synthetic system in cell culture should offer a unique opportunity to investigate the mechanisms of pattern formation and morphogen diffusion in detail. This work is also the first step to engineer a more complex synthetic tissue^1, 3-6^.

## Methods

### DNA constructs

The genetic constructs used in this study are listed in Supplementary table 1. The promoters or genes were subcloned into pDONR vector to create entry clones. These entry clones were recombined with *piggyBac* vector (a gift from Knut Woltjen)^44^ or Tol2 vector (a gift from Koichi Kawakami)^45,46^ by using the Multisite Gateway technology (Invitrogen). The (f2)_7_ promoter was created by fusing the (f2)_7_ enhancer sequence (a gift from Hiroshi Hamada)^33^ to the CMV minimal promoter. The CAG promoter is a gift from Junichi Miyazaki^47^. Genes related to Nodal signaling (Nodal, Lefty2, Cryptic, FoxH1 and Acvr2b) were cloned from mouse cDNA Mix (GenoStaff). CAAX domain was cloned from Kras4B. The CRISPR guide sequences for Acvr2b deletion were cloned into pSpCas9(BB)-2A-Puro vector (a gift from Feng Zhang, Addgene #62988)^48^.

### Cell culture

293AD (Cell Biolabs), a cell line derived from parental HEK293 cells, was used for all experiments because of its flattened morphology and firm attachment to a culture dish. The cells were maintained in DMEM/F12 medium containing 10% fetal bovine serum at 37 °C with 5% CO_2_.

### Creation of stable cell clones

The genetic constructs were introduced into HEK293 cells with the *piggyBac* or Tol2 transposase. To create the reporter cell line for Nodal signaling, (f2)_7_-luc, CAG-Cryptic, CAG-FoxH1 and CAG-Acvr2b were introduced into HEK293 cells. To create the activator circuit, (f2)_7_-Nodal was added to the reporter cell line. To create the activator-inhibitor circuit, (f2)_7_-Lefty2 was added to the activator cell line. To increase the maximum synthesis rate of Nodal or Lefty, (f2)_7_-Nodal with PGK-GFP or (f2)_7_-Lefty2 with PGK-mCherry was further added to the activator-inhibitor cell line. To create the similar range activator-inhibitor circuit, (f2)_7_-F1-Lefty2 was added to the activator cell line instead of (f2)_7_-Lefty2. To create HiBiT-tagged ligand cell lines, CAG-ligand and EF1a-mCherry-CAAX were introduced into HEK293 cells. After antibiotics selection, all cell lines except for the cells used in Fig 4f and Supplementary figs. 1 and 8b were cloned from a single cell.

### Time-lapse imaging of synthetic pattern formation

The 2.0×10^5^ cells (200 μl suspension) were seeded onto the glass part (φ12 mm) of a glass base 35 mm dish (IWAKI). After 2 hours incubation, the medium was replaced with the 2 ml fresh medium containing 20 mM HEPES and 100 μM D-luciferin, and the luminescence was imaged at each time point with a customized incubator microscope LCV110 (Olympus).

### Culture insert assay with the HiBiT system

A culture insert (Ibidi) was placed on the glass part of a glass base 35 mm dish, and then the ligand cells and receptor cells (4×10^4^ cells each) were seeded separately in the two wells of the culture insert. After 2 hours incubation, the culture insert was removed, and the cells were cultured with 2 ml medium. After 2-3 days incubation, the medium was replaced with the 2 ml fresh medium containing 20 mM HEPES, 1 μl HiBiT substrate (Nano-Glo^®^ Live Cell EX-4377, Promega) and 8 μl LgBiT (Promega), and the luminescence and fluorescence were imaged with LCV110. Cytochalasin B (5 μM) was added at 20 hours, and then measurement was resumed. Cells labeled with mCherry-CAAX were imaged with a confocal microscope LSM 780 (Carl Zeiss) in Supplementary fig. 6.

### Quantification of HiBiT activity

A 30×300 pixels (48×480 μm^2^) rectangular area was set so that the short side of the rectangle is in parallel with the boundary between the ligand cells and receptor cells. The HiBiT activities and mCherry intensities were averaged along the short side. The averaged mCherry intensities were normalized with the following function:

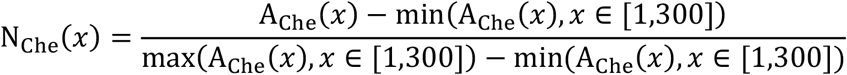

where A_Che_ (x) is the averaged mCherry intensity at position x pixel (x = 1 is the in the region of ligand cells and x = 300 is in the region of receptor cells). The averaged HiBiT activities were normalized with the following function:

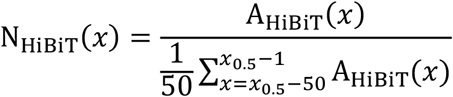

where A_HiBiT_(x) is the averaged HiBiT activity at position x, and x_0.5_ is the position where the normalized mCherry intensity drops to 0.5. The normalized HiBiT activities of three independent experiments were averaged and then fitted to the following function:

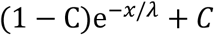

where C < 1 is background. Finally, the HiBiT activity distributions were given by

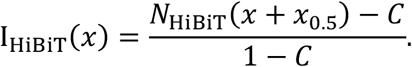

The distance was compensated by 1 pixel = 1.6 μm.

### Luciferase assay

For the luciferase assay shown in Supplementary fig. 1b, the cells were seeded in a 24- well plate at 1.0×10^5^ cells/well. After 24 hours culture in the absence or presence of 10 nM recombinant Nodal, the cells were washed with PBS and eluted with 150 μl 1×lysate buffer (Luciferase assay system, Promega). For the intermingled co-culture assay (Fig 4f; Supplementary fig. 2), the ligand cells and reporter cells were seeded in a 24-well plate at 1.0×10^5^ cells each/well and mixed. After 48 hours co-culture, the cells were washed with PBS and eluted with 250 μl 1×lysate buffer. For the measurement of the signal response curve (Supplementary fig. 8a), the reporter cells were seeded in a 96-well plate at 7000 cells/well. After 1 hour incubation, the medium was changed into the fresh medium containing recombinant Nodal and Lefty1. After 48 hours culture, the cells were washed with PBS and eluted with 100 μl 1×lysate buffer. The 20 μl lysate prepared above was mixed with 50 μl luciferase substrate (Luciferase assay system, Promega), and the luminescence was measured with a luminometer TriStar^2^ (Berthold technologies).

### Degradation rate

The 3×10^5^ cells expressing each HA-tagged ligand were seeded in a 35 mm dish. After 2 days culture, the cells were treated with cycloheximide (50 μg/ml) and sampled for immunoblotting at 0, 1, 2, 3 and 6 hours. Immunoblotting was performed according to a standard protocol, and the blot was probed with mouse anti-HA antibody (901501, Biolegend; 1/3,000). The resulting bands were quantified with an ImageJ plug-in Gel Analyzer.

### FACS analysis

The fluorescent cells were quantified with JSAN cell sorter (Bay bioscience) and analyzed with FlowJo software.

### Models

Two simple mathematical RD models of the activator-inhibitor circuit were constructed. In the competitive inhibition model, Lefty was assumed to inhibit Nodal by competing for the co-receptor and receptors.

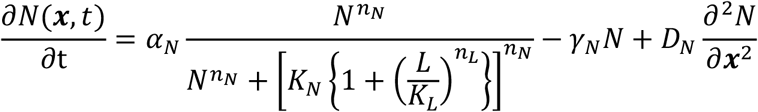

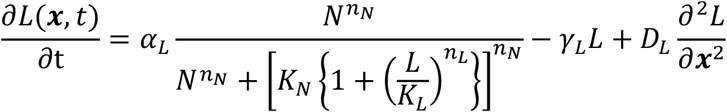

where N, α_N_, n_N_, K_N_, γ_N_ and D_N_ are the concentration, maximum synthesis rate, Hill coefficient, dissociation rate, degradation rate and diffusion coefficient of Nodal, respectively, and L, α_L_, n_L_, K_L_, γ_L_ and D_L_ are those of Lefty. In the competitive inhibition + direct inhibition model, Lefty was also assumed to inhibit Nodal by directly binding to it.

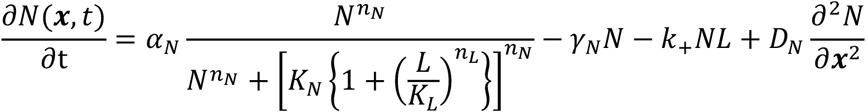

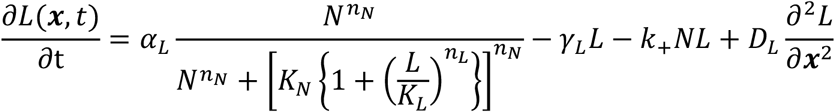

where k_+_ is the binding rate constant of Nodal and Lefty.

Numerical simulations of the models were performed by using simple Euler method or ode45 solver of MATLAB (Mathworks). The diffusion terms were numerically solved by using the finite difference method. Parameters used are n_N_ = 2.63, K_N_ = 9.28 nM, γ_N_ = 2.37×10^-3^ min^-1^, D_N_ = 1.96 μm^2^/min, n_L_ = 1.09, K_L_ = 14.96 nM, γ_L_ = 5.65×10^-3^ min^-1^, D_L_ = 56.39 μm^2^/min and k_+_ = 0.03 min^-1^nM^-1^, unless stated otherwise.

To create the phase diagram of pattern forming ability, 1D simulations with different combinations of α_N_ and α_L_ were performed for much longer time than the time scale of our experiment. As the initial state, two pulses of Nodal and Lefty concentrations were set. The resulting Nodal distribution was judged as a “pattern” when the maximum Nodal concentration was more than 2-times higher than the minimum Nodal concentration. Otherwise, the distribution was judged as a “high state” or “low state”, depending on if the maximum Nodal concentration was higher than 0.01 or not.

The Turing instability condition was judged based on the following four inequalities^13^:

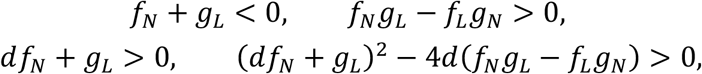

where *f_N_, f_L_* are the derivatives of *f* with respect to *N*, *L,* respectively, and *g_N_, g_L_* are the derivatives of *g* with respect to *N*, *L*. Functions *f* and *g* are given by

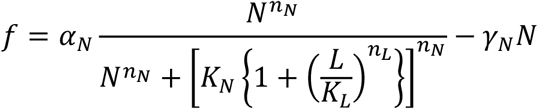

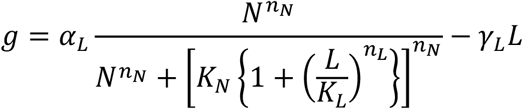

for the competitive inhibition model, or

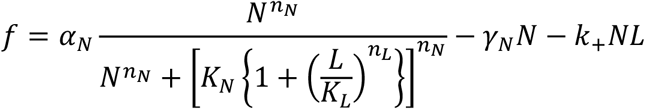

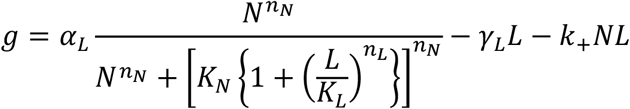

for the competitive inhibition + direct inhibition model.

### Spatial correlation analysis

The correlation function of the florescent intensity is given by

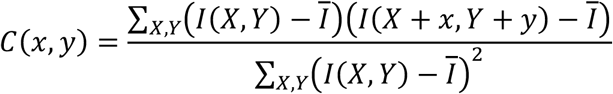

where the summation was taken over all pixel points (*X,Y*) and *I*̄ is the mean intensity. The radial correlation function *C*(*r*) was calculated by averaging the correlation function *C*(*x,y*) with the constraint 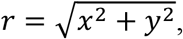 which is formally given by

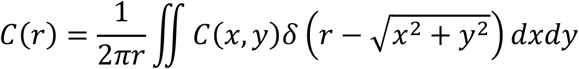

Where *δ*(*x*) is the Dirac’s delta function.

## Acknowledgments

We thank M. Matsuda for helping experiments, X. Diego and J. Sharpe for helping data analyses, H. Hamada for helpful advice and the members of Ebisuya lab for technical assistance and discussion. This work was supported by Precursory Research for Embryonic Science and Technology (PRESTO) (JPMJPR12A6 to M.E.), Grant-in-Aid for Scientific research (KAKENHI) programs from Ministry of Education Culture, Sports, Science, and Technology (MEXT) (16KT0080 to M.E., 26891027 to R.S.) and RIKEN Special Postdoctoral Researchers (SPDR) fellowship to R.S..

## Author contributions

R.S. and M.E. designed the work and wrote the manuscript. R.S. performed the experiments. R.S. and T.S. constructed the models and analyzed the data.

---

**Supplementary figure 1.**
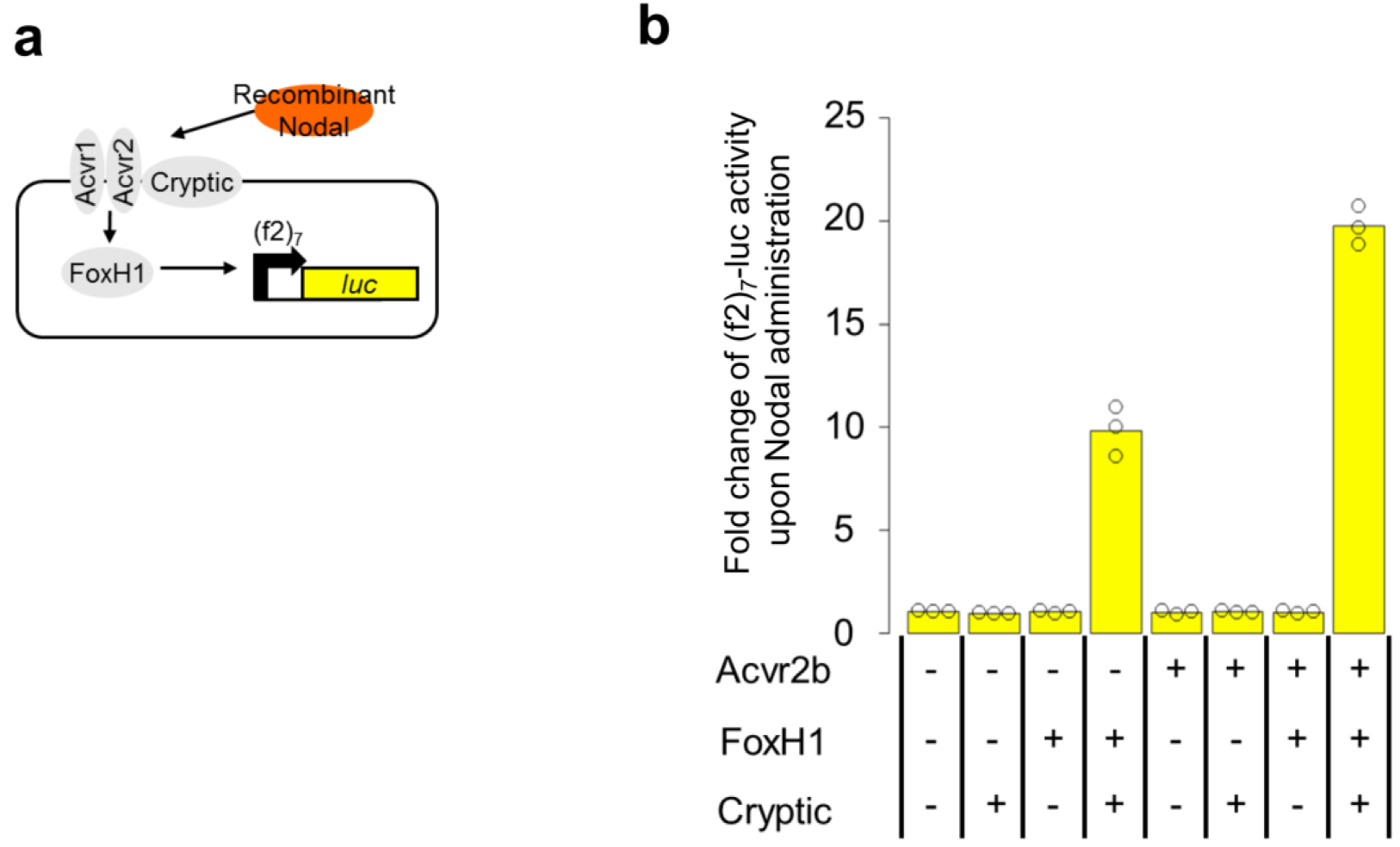
Cryptic, FoxH1 and Acvr2b are required for HEK293 cells to efficiently respond to Nodal stimulation. (**a**) Different combinations of Cryptic (co-receptor), FoxH1 (transcription factor) and Acvr2b (receptor) were introduced into HEK293 cells, and the Nodal signaling activities were monitored with the (f2)7-luc reporter. (**b**) The (f2)7-luc activities in the absence and presence of 10 nM recombinant Nodal were measured, and the fold-changes are shown. Data are means and individual points (n=3).

**Supplementary figure 2.**
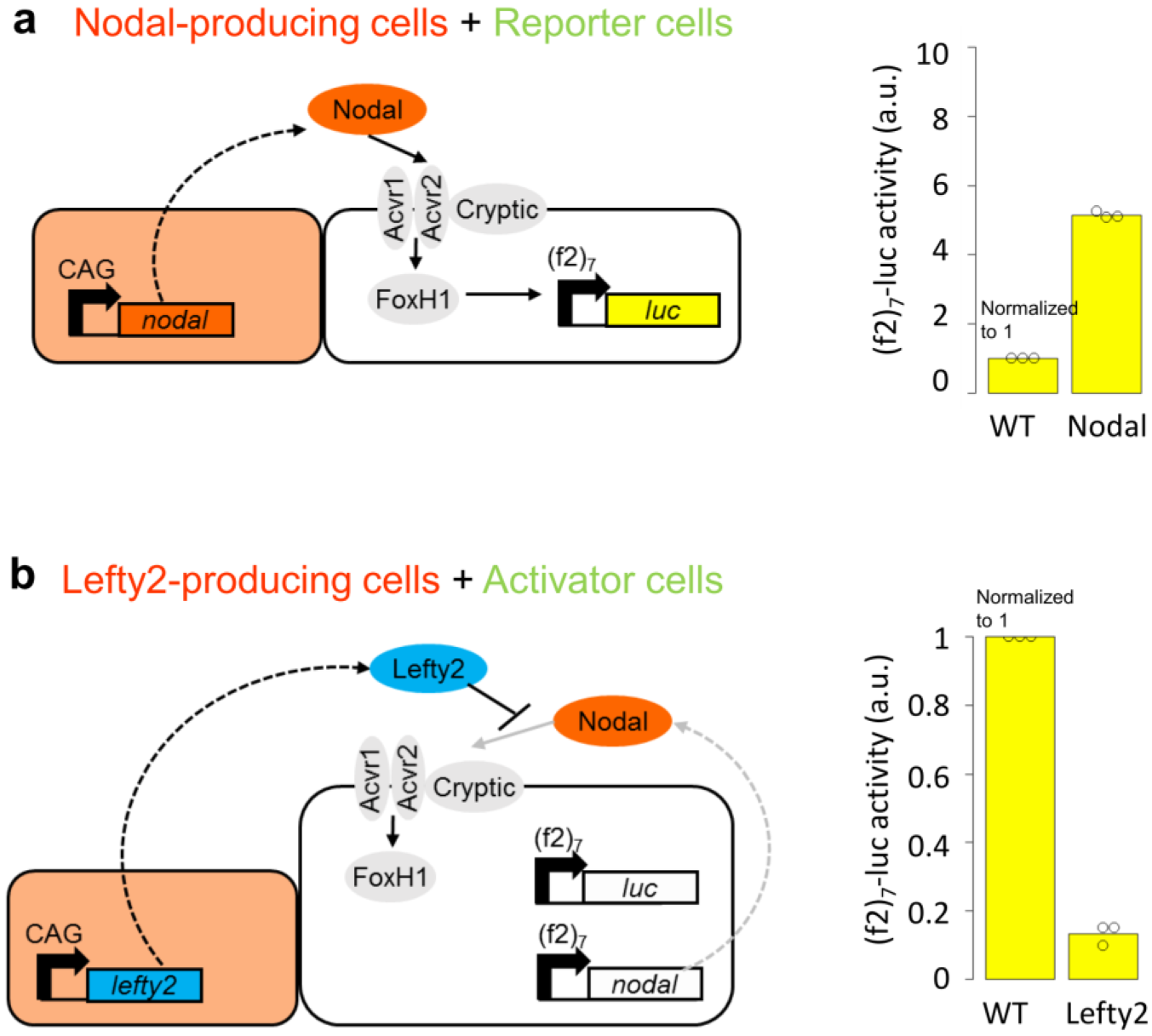
Co-culture with Nodal-or Lefty2-producing cells. (**a**) The Nodal-producing cells or wild-type cells were co-cultured with the reporter cells, and the (f2)_7_-luc activity of the reporter cells was measured 48 hours later. (**b**) The Lefty2-producing cells or wild-type cells were co-cultured with the activator cells, and the (f2)_7_- luc activity of the activator cells was measured 48 hours later. Data are means and individual points (n=3).

**Supplementary figure 3.**
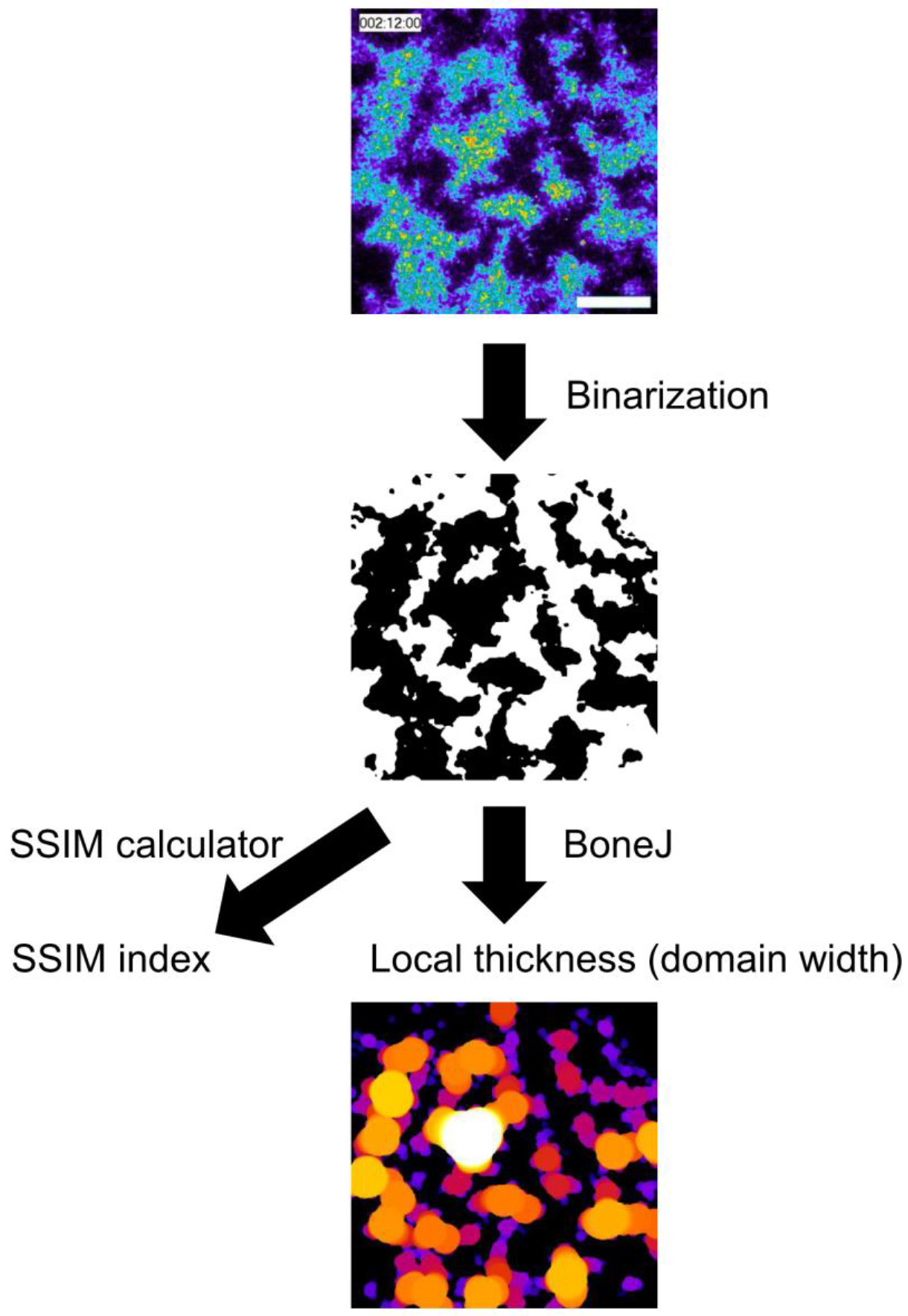
A scheme of pattern analyses. Raw luciferase images were binarized (threshold = the median of luminescence distribution). Then the structure similarity (SSIM) index and local thickness were calculated with the ImageJ plugins, SSIM_INDEX and BoneJ, respectively. The average positive/negative domain width was defined as the average value of the local thickness.

**Supplementary figure 4.**
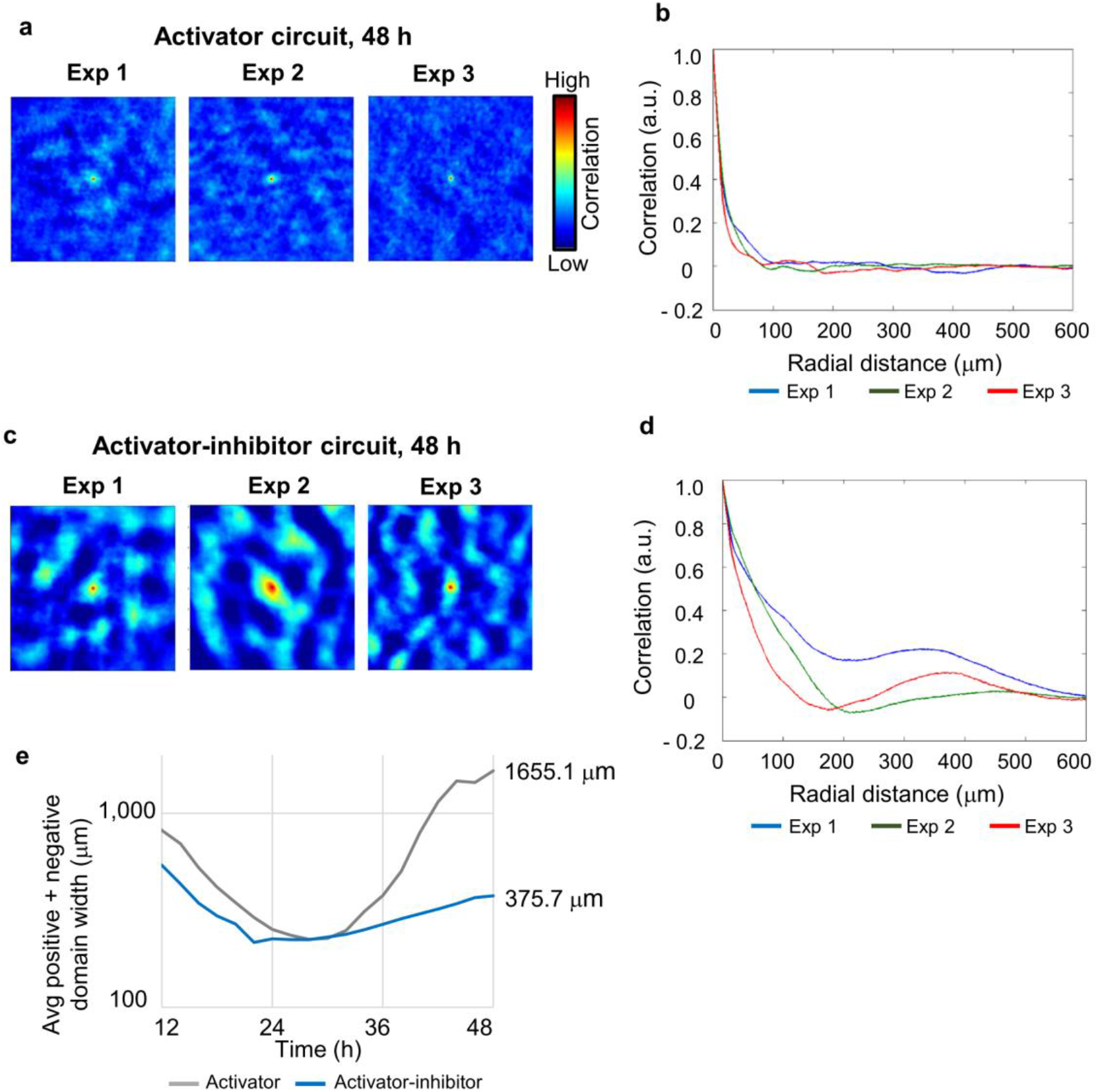
Spatial correlation analyses of the synthetic pattern. (**a**) The spatial correlation was calculated for the image of the cells engineered with the activator circuit. (**b**) The correlation function shown in (a) was radially averaged and plotted. (**c**) The spatial correlation was calculated for the pattern of the cells engineered with the activator-inhibitor circuit. (**d**) The correlation function shown in (c) was radially averaged and plotted, revealing a second peak at around 400 μm. (**e**) The sum of the average positive domain width and negative domain width at each time point.

**Supplementary figure 5.**
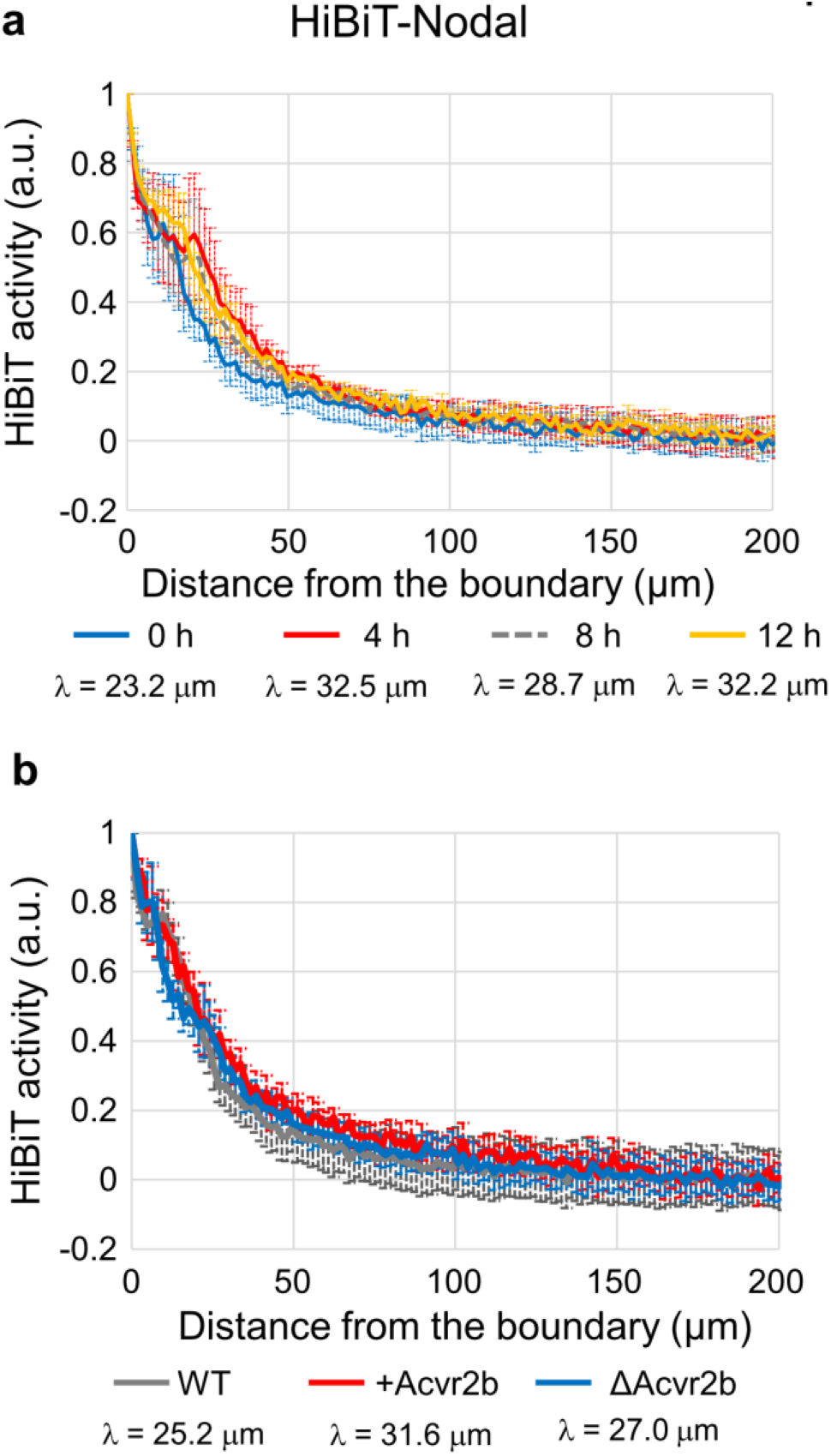
Extracellular distribution of Nodal. (**a**) Quantified distribution profiles of HiBiT-Nodal at 0, 4, 8 and 12 hours after the addition of the LgBiT and substrate in the culture insert assay. The data at 8 hours is also shown in Fig. 2e. (**b**) The effect of Acvr2b on the distribution range of HiBiT-Nodal. HiBiT-Nodal was introduced into the wild-type cells, Acvr2b-overexpressing cells and Acvr2b-knock out cells. These ligand cells were co-cultured with the corresponding receptor cells (wild-type cells, Acvr2b-overexpressing cells and Acvr2b-knock out cells, respectively) in the culture insert assay.

**Supplementary figure 6.**
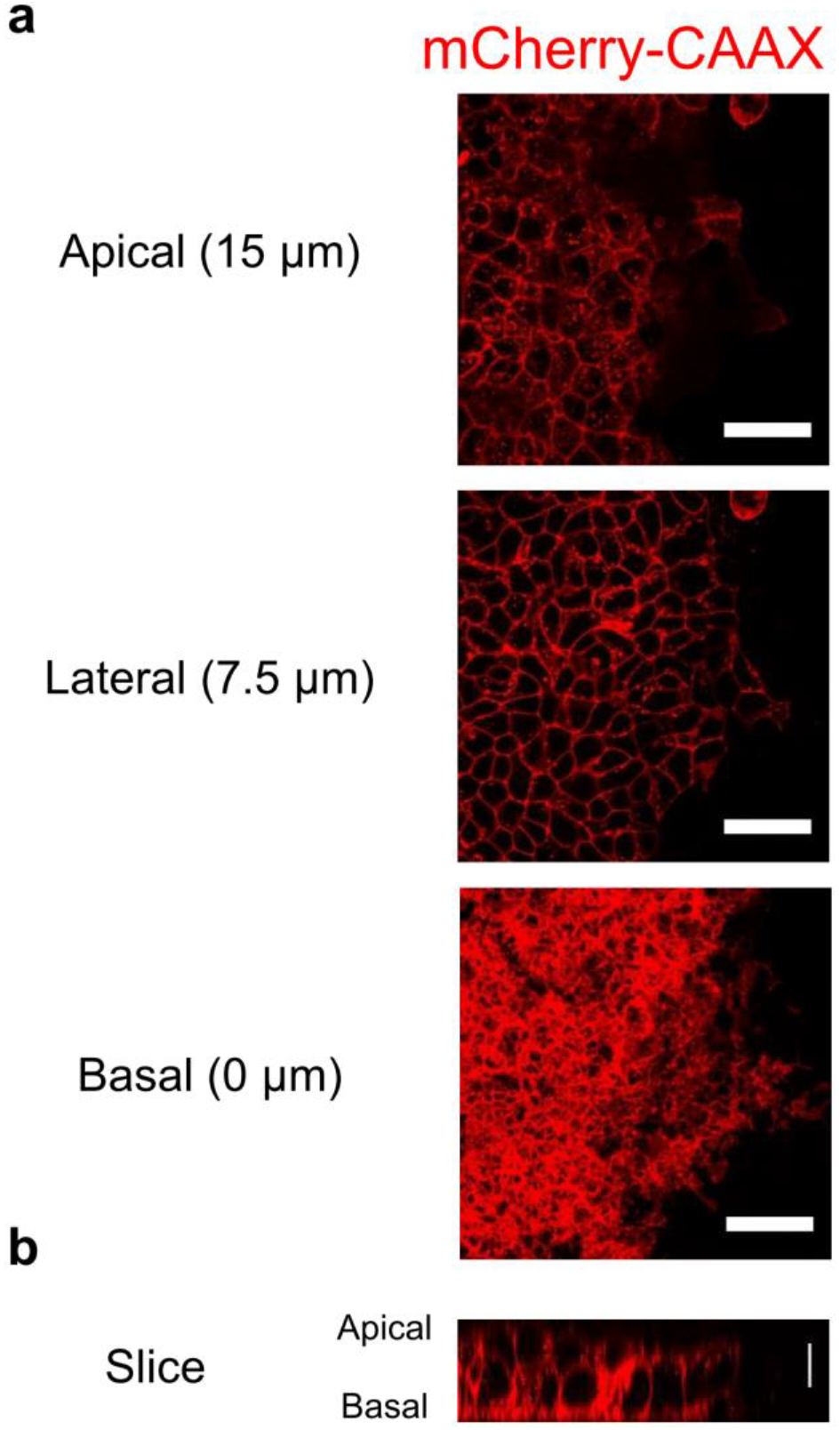
A z-stack view of ligand-producing cells. (**a,b**) The ligand cells (labeled with mCherry-CAAX) in the culture insert assay were imaged with a confocal microscope. The basal side was defined as the point where the cell membranes showed a dense structure spreading on the dish. The lateral side and apical side were defined as the points 7.5 μm and 15 μm higher than the basal side, respectively. Scale bars, 50 μm (a); 10 μm (b).

**Supplementary figure 7.**
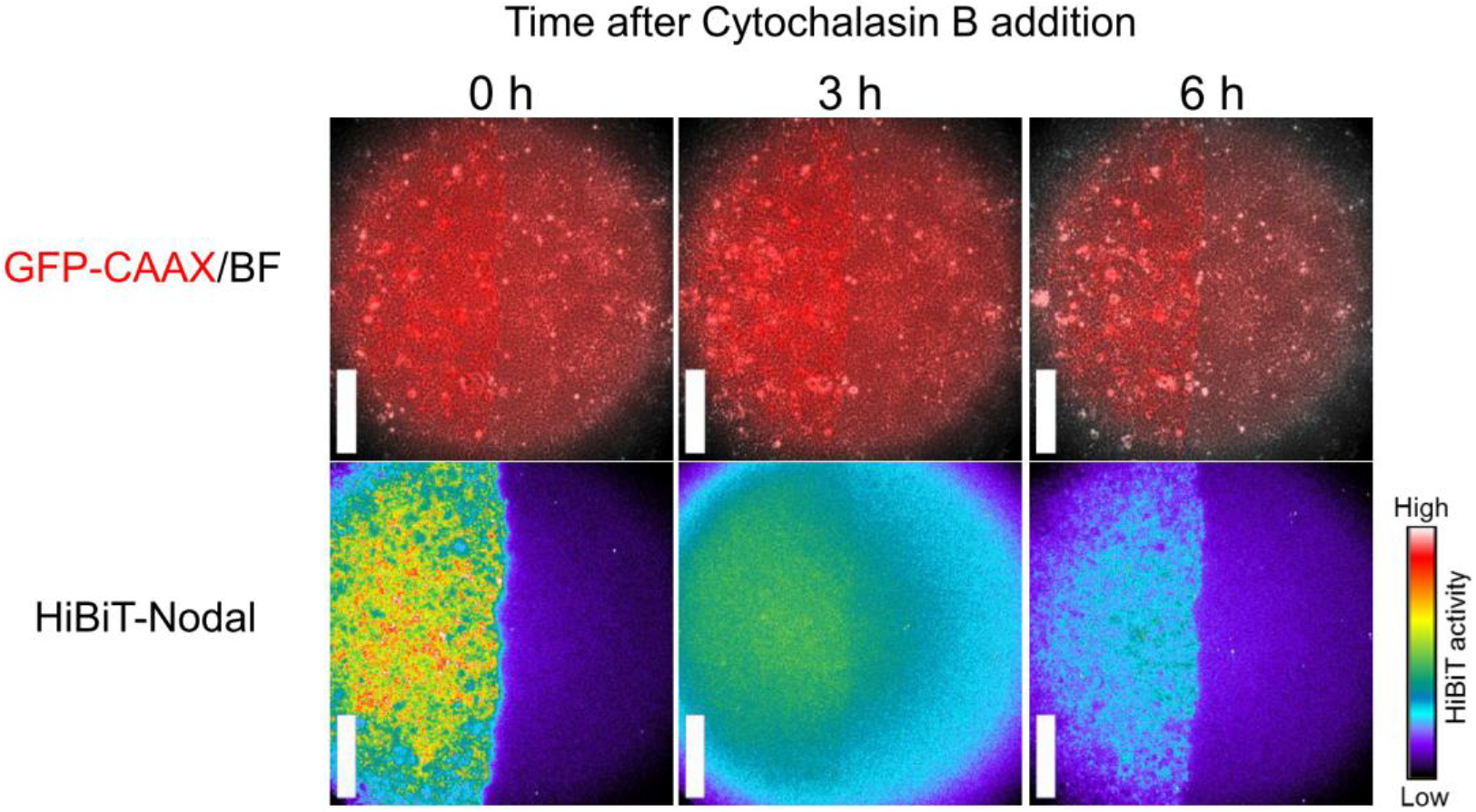
The effect of Actin inhibition on the Nodal distribution. The HiBiT-Nodal signal was imaged at 0, 3 and 6 hours after the addition of Cytochalasin B (5 μM) in the culture insert assay. The ligand cells were labeled with GFP-CAAX. Scale bars, 400 μm.

**Supplementary figure 8.**
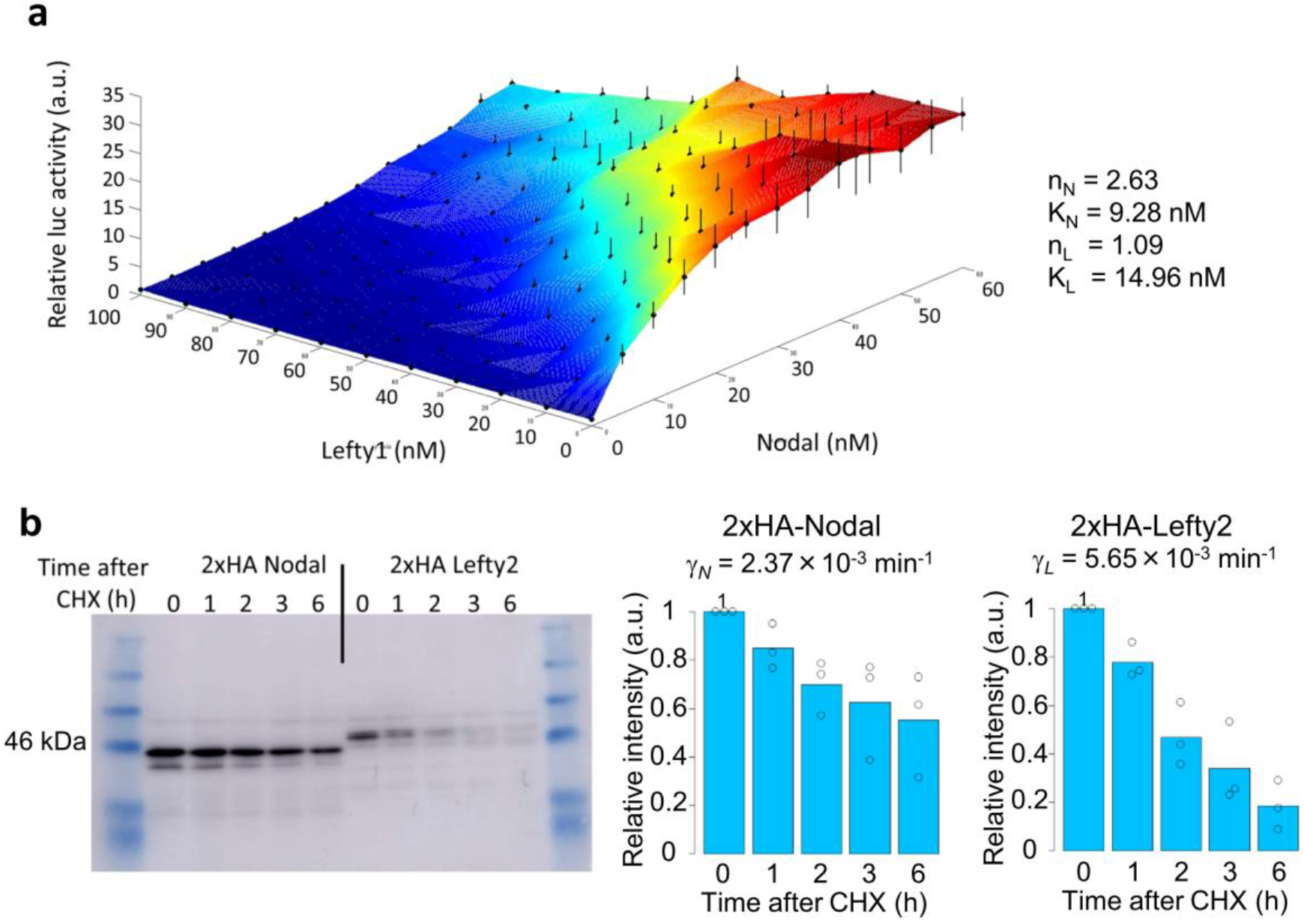
Parameter measurements. (**a**) Measurements of the signal response curve. The reporter cells containing (f2)_7_-luc, Cryptic and FoxH1 were treated with different concentrations of recombinant Nodal and Lefty1. The (f2)_7_-luc activities were measured 2 days later. Data are means and s.e.m. (n=3). (**b**) Measurements of the degradation rates. The protein amounts of the HA-tagged Nodal (43.7 kDa) and HA-tagged Lefty2 (44.6 kDa) were measured at each time point after the addition of cycloheximide (CHX, 50 μg/ml). Left: A representative gel image of the immunoblotting. Center and right: The quantification of (b). Data are means and individual points (n=3). Note that the extracellular and intracellular proteins were not distinguished in this measurement. Using the degradation rates γ and the characteristic distances λ measured in Fig. 2e, the diffusion rates D of Nodal and Lefty2 were estimated as DN = 1.96 μm^2^/min and DL = 56.39 μm^2^/min.

**Supplementary figure 9.**
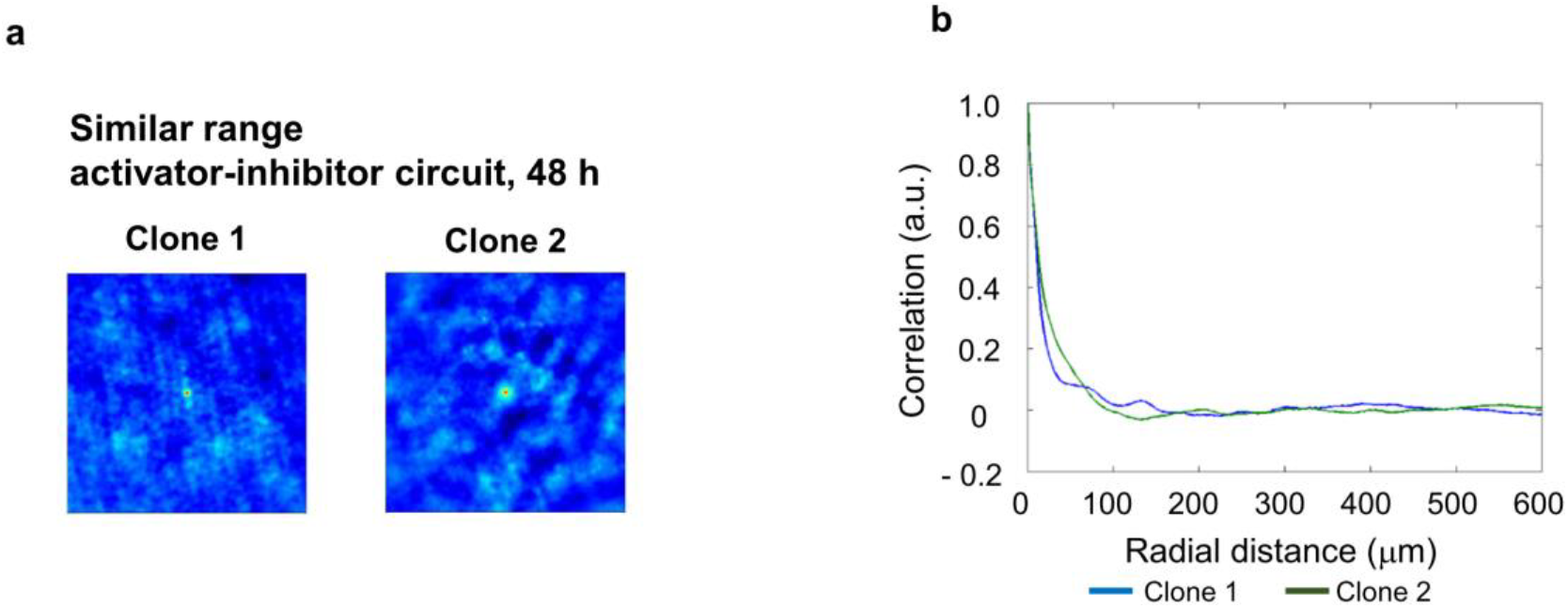
Spatial correlation analyses of the similar range activator-inhibitor circuit. (**a**) The spatial correlation was calculated for the image of the cells engineered with the similar range activator-inhibitor circuit. (**b**) The correlation function shown in (a) was radially averaged and plotted.

**Supplementary movie 1.**

The time-lapse imaging of the HEK293 cells engineered with the activator circuit and that with the activator-inhibitor circuit. Scale bar, 400 μm.

